# Sublethal insecticide exposure of larvae affects the blood-feeding behaviour of adult mosquitoes

**DOI:** 10.1101/2024.06.26.600778

**Authors:** Tiago G. Zeferino, Gwendoline Acerbi, Jacob C. Koella

**Affiliations:** Institute of Biology, University of Neuchâtel, Rue Emilie-Argand 11, 2000 Neuchâtel, Switzerland

## Abstract

Because of their widespread use for the control of disease vectors and agricultural pests, insecticides have become ubiquitous in the environment, including in water bodies harbouring mosquito larvae. These are therefore continuously exposed to sublethal doses. Since this has long-lasting effects on the mosquitoes’ physiology and life-history, we expected that it may also affect behaviours that underlie the mosquitoes’ population dynamics and disease epidemiology, such as egg-laying preference, blood-feeding motivation, and host-seeking behaviour. Using an insecticide-sensitive and a resistant strain of *Anopheles gambiae*, an important malaria vector, we evaluated the effects of sublethal exposure to permethrin throughout larval development on the resistance to the insecticide in adults, on host-seeking behaviour, on the motivation to blood-feed, and on egg-laying behaviour. Resistance, assessed by rates of knock-down and mortality, were similar between exposed and unexposed mosquitoes. However, exposure to sublethal doses of insecticide caused female mosquitoes to split their egg clutches into two parts and increased the motivation of mosquitoes to seek blood meals through permethrin-treated nets, regardless of their sensitivity to the insecticide. Furthermore, it enhanced the natural preference of resistant strains for permethrin-treated nets and increased their blood-meal size. Our results thus suggest that sublethal insecticide concentrations in larval breeding sites have important epidemiological implications.

## Introduction

Mosquitoes and other vectors of infectious diseases are often exposed to sublethal doses of insecticides. One reason is that these chemicals are extensively used to control mosquito-borne diseases like malaria, in particular in combination with bed-nets ^1,2^. Therefore, in searching for a host, mosquitoes will touch a bed-net and can be irritated by the insecticide and fly away before the contact is long enough to kill them. Another reason is that some of the insecticides that are used in agriculture often end up in surface or ground water ^3,4^, through various pathways such as atmospheric deposition, soil leaching ^5^, and volatilization or degradation ^6^, where they can affect many animals, including aquatic arthropods such as mosquito larvae.

Even if their concentration is too low to kill the larvae, these insecticides impact the adults by influencing their life-history traits ^6–9^, their behaviour ^10–12^, and their competence to transmit, for example, arboviruses ^13,14^ and malaria parasites ^15–17^. Mechanisms can be that insecticides interfere with important physiological pathways ^18^. Insecticides, for example, impair the olfactory systems of mosquitoes ^19,20^, thus impeding their responses to attractants involved in host-seeking and oviposition ^11,21–23^. Some effects of an exposure to a low dose of insecticide can carry on to the next generation: the offspring of mothers that had been exposed to a low dose of a pyrethroid were bigger as larvae and pupae than those of unexposed individuals ^24^, while a low dose of spirotetramat decreased body size and longevity in the offspring of cabbage aphids ^25^.

Despite this body of knowledge, little is known about how the exposure to a sublethal dose of insecticide affects the later response of mosquitoes to insecticides. One exception is an experiment suggesting that repeated exposure to permethrin in adults does not change sensitivity to a subsequent exposure ^26^.

That the stress of exposure affects the response to insecticides would not be too surprising, for other stressors do the same. Malaria-infected mosquitoes, for example, are less repelled by insecticide-treated nets suggesting that they could also be less killed ^27^. Larval crowding ^28^ or undernutrition ^29^ both also increase the phenotypic sensitivity of mosquitoes to insecticides.

In several experiments, we investigated how sublethal larval exposure to permethrin influences adult mosquitoes’ responses to insecticide in both an insecticide-resistant and a sensitive strain of *Anopheles gambiae*. We assessed resistance by measuring knock-down rates and mortality using the WHO tube test. Additionally, we investigated the mosquitoes’ preference for a host protected by an insecticide-treated or untreated net, their motivation and ability to bite through a permethrin-treated net, and their choice of insecticide-laced or untreated water as an oviposition site.

## Results

### 1. Insecticide resistance

We exposed adult mosquitoes (690 Kisumu-mosquitoes and 507 RSP-mosquitoes) to 0.75% permethrin-impregnated paper in WHO test tubes for 30 or 60 minutes, and assessed whether they were knocked down during the exposure and whether they died within 24 hours after the exposure (**Fig. 5c**).

Kisumu-mosquitoes were knocked down faster than RSP (median knock-down times: 15 minutes vs 60 minutes, χ^2^ = 531.24, df = 1, p < 0.001), but the rate at which mosquitoes were knocked-down was not affected by either larval exposure to permethrin (χ^2^ = 0.27, df = 1, p = 0.605) or the interaction between larval exposure and strain (χ^2^ = 0.34, df = 1, p = 0.561, **Fig. 1a**). In addition, almost all of the Kisumu-mosquitoes were knocked down after 30 minutes (94% vs 100% after 60 minutes), while of the RSP-mosquitoes only 11% were knocked down after 30 minutes but 80% were knocked down after 60 minutes (interaction strain * duration of exposure: χ^2^ = 20.64, df = 1, p < 0.001).

**Figure 1.**
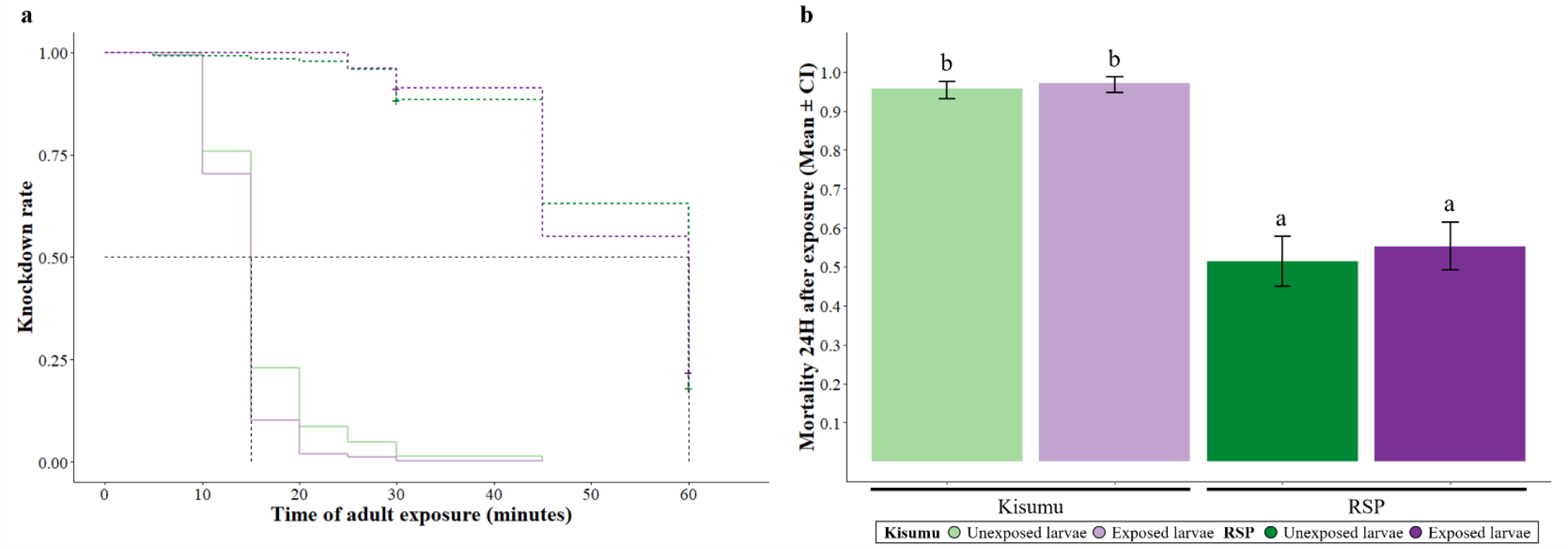
Insecticide resistance. The proportion of individuals **(a)** knockdown or **(b)** dead 24 hours after their adult exposure for each strain and larval exposure treatment. Letters indicate statistically significant differences from multiple comparisons. For details on the sample size see **Supplementary Table S1.2**.

Almost all of the Kisumu-mosquitoes (96%) died within 24 hours of exposure, while only 53% of RSP died (χ^2^ = 180.83, df = 1, p < 0.001). In addition, among Kisumu-mosquitoes there was close to 100% mortality already after 30 minutes, while among RSP-mosquitoes mortality increased from 31% after 30 minutes of exposure to 75% after 60 minutes (interaction strain * duration of exposure: χ^2^ = 20.65, df = 1, p < 0.001). Mortality was not affected neither by larval exposure to permethrin (78% for unexposed and for exposed mosquitoes, χ^2^ = 2.34, df = 1, p = 0.125) nor by the interaction between larval exposure and strain (χ^2^ = 0.34, df = 1, p = 0.561, **Fig. 1b**). For further details on the statistical analysis see **Supplementary Table S1.1**.

### 2. Choice of protected or unprotected host

Placed into a central cage, females were given the opportunity to move into a cage where its blood-meal source (one of our arms) was protected by a permethrin-treated net or by an untreated net (**Fig. 5d**).

Of the 687 mosquitoes, about 32% left the central cage, independently of treatment (strain: χ^2^ = 1.41, df = 1, p = 0.234; larval exposure: χ^2^ = 0.97, df = 1, p = 0.322; interaction strain * larval exposure: χ^2^ = 3.58, df = 1, p = 0.058, **Fig. 2a**).

**Figure 2.**
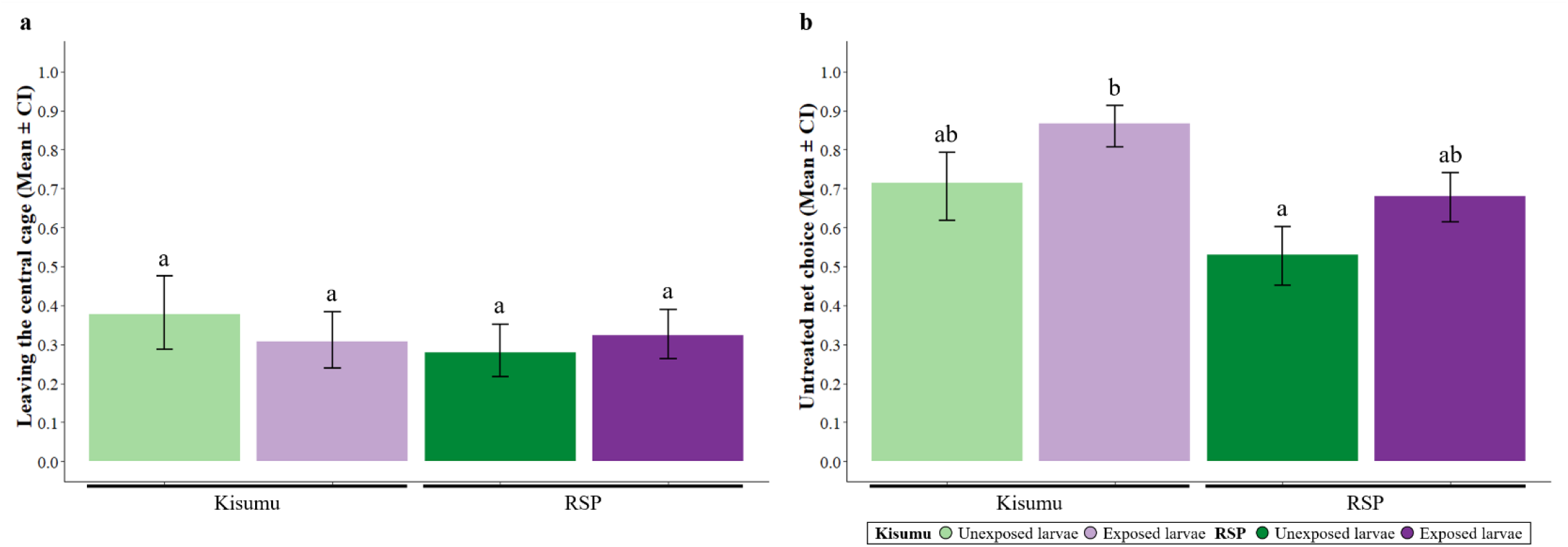
Choice of protected or unprotected host. The proportion of individuals that **(a)** left the central cage and **(b)** chose the untreated net side for each strain and larval exposure. Letters indicate statistically significant differences from multiple comparisons. For details on the sample size see **Supplementary Table S2.2**.

Among the 218 mosquitoes that did leave the central cage, Kisumu-mosquitoes (80%) were more likely to choose the untreated net than RSP-mosquitoes (61%) (χ^2^ = 4.22, df = 1, p = 0.039), and mosquitoes exposed as larvae (76%) were more likely than unexposed ones (61%) to choose the untreated net (χ^2^ = 6.08, df = 1, p = 0.013), independently of them being Kisumu-mosquitoes (86% for exposed vs 71% for unexposed) or RSP-mosquitoes (68% for exposed vs 52% for unexposed, interaction strain * exposure: χ^2^ = 0.23, df = 1, p = 0.626, **Fig. 2b**). For further details on the statistical analysis see **Supplementary Table S2.1**.

### 3. Motivation and ability to bite through a permethrin-treated net

Females that had been exposed or not exposed as larvae were allowed to blood-feed for five minutes through a permethrin-treated or an untreated net (**Fig. 5e**).

Of the 582 mosquitoes, 57% of Kisumu-mosquitoes and 42% of RSP-mosquitoes took a blood meal (χ^2^ = 13.84, df = 1, p < 0.001). While the motivation of Kisumu-mosquitoes to take a blood-meal was only slightly affected by the type of net (60% bit through an untreated net and 54% through a treated next), RSP-mosquitoes were more likely to take a blood-meal through a permethrin-treated net (50%) than through an untreated net (35%) (interaction strain * net: χ^2^ = 5.79, df = 1, p = 0.016). Overall, the proportion that took a blood meal was not affected by larval exposure (χ^2^ = 0.92, df = 1, p = 0.335) or by the treatment of the net (χ^2^ = 0.91, df = 1, p = 0.337). However, exposed individuals were more likely to take a blood-meal through a permethrin-treated net (58%) than through an untreated net (45%), whereas unexposed individuals tended to avoid biting through the treated net (45% for permethrin-treated net vs. 50% for untreated net; interaction exposure * net: χ^2^ = 4.74, df = 1, p = 0.029, **Fig. 3a**).

**Figure 3.**
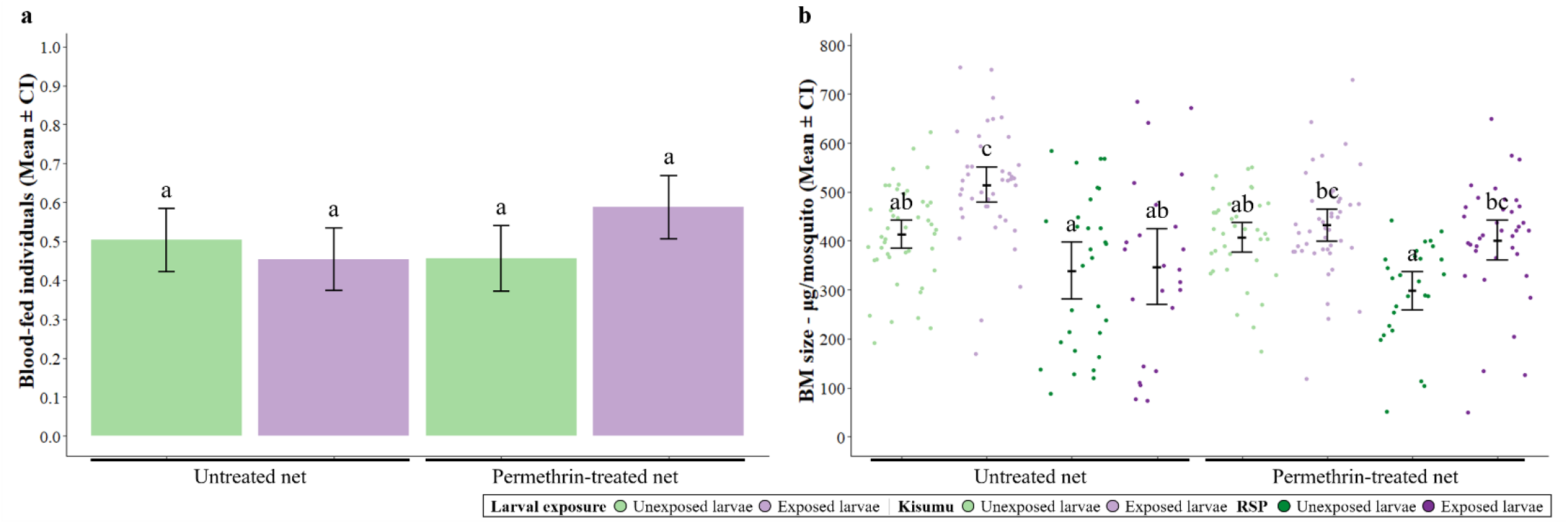
Motivation and ability to bite through a permethrin-treated net. **(a)** The proportion of individuals that took a blood-meal through a permethrin-treated or untreated net and **(b)** their individual blood-meal size (i.e., quantity of haemoglobin) for each larval exposure treatment and strain. Letters indicate statistically significant differences from multiple comparisons. For details on the sample size see **Supplementary Table S3.2**.

Among the mosquitoes that took a blood-meal, Kisumu-mosquitoes took up more blood (467.43 μg/mosquito) than RSP-mosquitoes (357.85 μg/mosquito) (F_1,283_ = 3.54, df = 1, p = 0.060). Mosquitoes exposed as larvae took larger bloodmeals (467.36 μg/mosquito) than unexposed mosquitoes (373.02 μg/mosquito) (F_1,283_ = 17.93, df = 1, p < 0.001), in particular in Kisumu-mosquitoes (interaction exposure * strain: F_1,283_ = 5.99, df = 1, p = 0.014). Overall, mosquitoes took up similar amounts of blood through treated (411.69 μg/mosquito) and untreated (433.78 μg/mosquito) nets (F_1,283_ = 0.03, df = 1, p = 0.852), but the difference of the bloodmeal size between treated and untreated nets depended on the combination of the strain and larval exposure (interaction exposure * strain * net: F_1,283_ = 6.04, df = 1, p = 0.014) (**Fig. 3b**). For further details on the statistical analysis see **Supplementary Table S3.1**.

### 4. Choice of egg-laying site

After mating and blood-feeding, females were given the choice between insecticide-laced and untreated water as an oviposition site (**Fig. 5f**).

Of the 353 females, 19% did not lay eggs. While this proportion was higher in RSP-mosquitoes (29%) than in Kisumu-mosquitoes (9%) (χ^2^ = 22.58, df = 1, p < 0.001), it was not affected by larval exposure to permethrin (χ^2^ = 0.02, df = 1, p = 0.882) or the interaction between larval exposure and strain (χ^2^ = 0.76, df = 1, p = 0.380, **Fig. 4a**). The average number of eggs was 76, with differences between treatments reflecting those for egg-laying likelihood (strain: F_1,281_ = 58.09, df = 1, p < 0.001; exposure: F_1,281_ = 0.93, df = 1, p = 0.333; interaction exposure * strain: F_1,281_ = 3.51, df = 1, p = 0.061, **Fig. 4b**).

**Figure 4.**
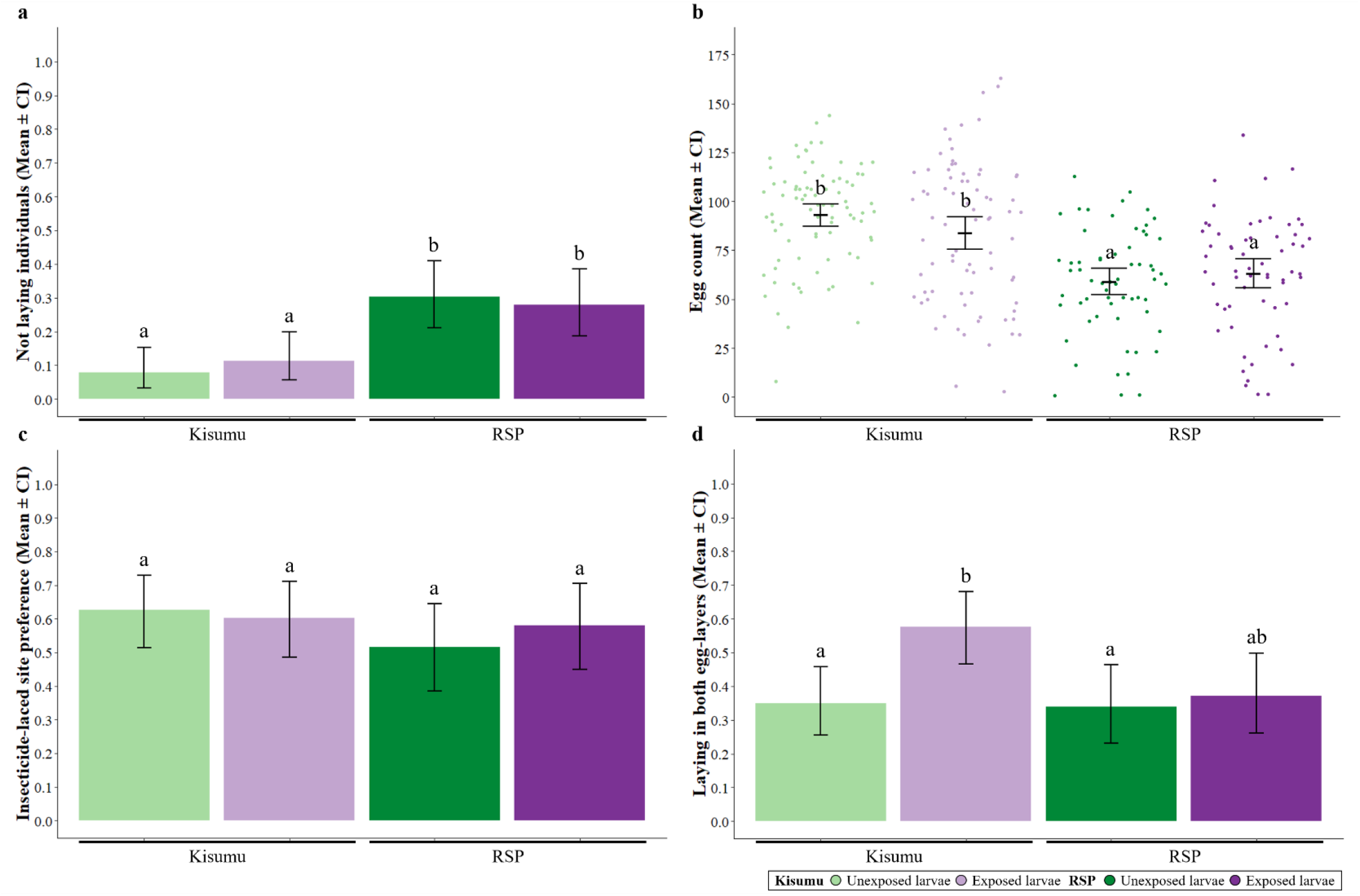
Choice of egg-laying site. The proportion of Kisumu and RSP individuals, exposed or unexposed to permethrin during the larval stage that **(a)** did not lay eggs or **(d)** laid in both egg-layers, as well as **(b)** the total egg count for each individual and **(c)** the proportion of eggs laid in the insecticide-laced site. Letters indicate statistically significant differences from multiple comparisons. For details on the sample size see **Supplementary Table S4.2**.

**Figure 5.**
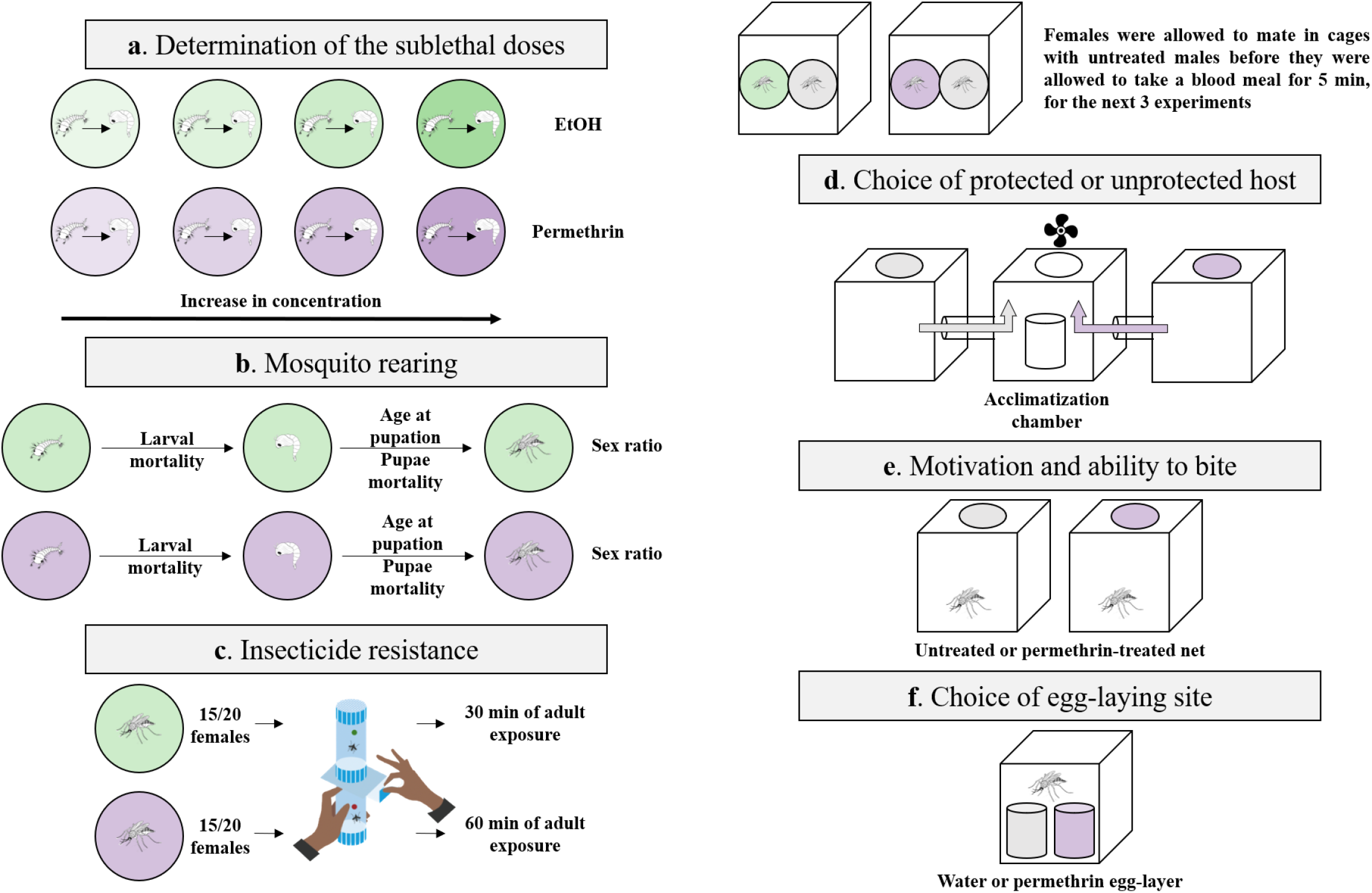
Experimental design. Illustration of the experiments in this study. **(a)** Determination of the sublethal doses, **(b)** Mosquito rearing, **(c)** Insecticide resistance, **(d)** Choice of protected or unprotected host, adapted from ^27^, **(e)** Motivation and ability to bite through a permethrin-treated net, **(f)** Choice of egg-laying site.

On average, 57% of eggs were laid into the insecticide-laced cup, with no effect of strain (χ^2^ = 1.28, df = 1, p = 257), larval exposure to permethrin (χ^2^ = 0.06, df = 1, p = 0.882) or the interaction between the two (χ^2^ = 0.55, df = 1, p = 0.455, **Fig. 4c**).

Of the 285 females that did lay eggs, 41 % laid into both egg-layers. This proportion was higher for exposed (48%) than for unexposed mosquitoes (34%) (χ^2^ = 6.05, df = 1, p = 0.013), in particular for Kisumu-mosquitoes (**Fig. 4d**). Kisumu-mosquitoes were slightly more likely to lay into both cups (χ^2^ = 3.39, df = 1, p = 0.065) in particular if they had been exposed, though the interaction between strain and exposure was not statistically significant (χ^2^ = 2.53, df = 1, p = 0.111). For further details on the statistical analysis see **Supplementary Table S4.1**.

## Discussion

Larval exposure to sublethal doses of insecticides significantly affected the physiology and behaviour of adult mosquitoes. Notably, while unexposed mosquitoes avoided biting through a permethrin-treated net, mosquitoes exposed during the larval stage preferred untreated nets when given a choice between a host protected by a treated and an untreated net (**Fig. 2**). However, when they were given no choice, exposed mosquitoes were more likely to take a blood meal through a treated net (**Fig. 3**). Exposed individuals exhibited increased skip oviposition behaviour (i.e. laying eggs in both available sites, **Fig. 4**). However, resistance, assessed as knockdown and mortality rates, was not affected by larval exposure (**Fig. 1**).

In contrast to other larval stressors like nutrition ^29^ and parasitism ^27^, larval exposure to permethrin did not affect insecticide resistance of the susceptible Kisumu or the resistant RSP-mosquitoes (**Fig. 1**). Although larval exposure influences oxidative homeostasis ^30^, which is crucial for resistance, our results align with previous studies indicating no impact on mortality from subsequent insecticide exposure ^26^.

When given a choice between a host protected by a treated or an untreated net (**Fig. 2b**), Kisumu-mosquitoes avoided the treated one, highlighting their inherent sensitivity to insecticides ^31^. Conversely, RSP-mosquitoes did not show this avoidance, likely due to the *kdr* mutation, which reduces the efficacy of insecticide-targeting voltage-gated sodium channels and alters the neural circuits, and therefore affects the mosquitoes’ sensitivity and the repellence of environmental cues ^22,23^. The difference between Kisumu and RSP corroborates several studies showing that resistant mosquitoes typically are repelled less than sensitive ones ^32–34^. Sublethal larval exposure to permethrin increased the repellency of the insecticide to the adults of both strains. A mechanism for this response would be that pyrethroid exposure affects the neural system in insects ^35^, altering their response to olfactory cues ^19,20^. Our results thus suggest that this effect is long-term, lasting from the exposure in larvae to the adult stage. Larval exposure thus sensitizes mosquitoes in a way that enhances their avoidance behaviour (i.e. excito-repellency ^36,37^) as adults, even if this behaviour is normally suppressed by a resistant mutation.

When allowed to bite either through an untreated or a treated net, but not given a choice between the two (**Fig. 3**), Kisumu-mosquitoes bit through the two nets with a similar likelihood. In contrast, RSP-mosquitoes were more likely to bite through a treated net, corroborating several studies ^7,30,38–40^. Furthermore, unexposed individuals bit through the two nets with a similar likelihood, but exposed individuals were more likely to bite through a permethrin-treated net than through an untreated one. While this appears to contradict our idea of sensitization, larval exposure is also a stress factor that might compel mosquitoes to prioritize fecundity, thus enhancing the motivation to bite. This behaviour aligns with the natural preferences of each strain, with Kisumu-mosquitoes favouring untreated nets and RSP-mosquitoes favouring treated nets (**Fig. 3b**). Whatever the mechanism, our results indicate a complex interplay of stress responses, strain-specific adaptations, and potential changes in sensory perception or behavioural conditioning due to larval insecticide exposure.

As in many studies of *Anopheles* mosquitoes ^41–43^, the resistant RSP-mosquitoes laid fewer eggs than the sensitive Kisumu-mosquitoes (**Fig. 4ab**). This suggests a cost of resistance, paid by re-allocating resources away from fecundity and towards resistance. In contrast to our prediction, larval exposure did not make females avoid the insecticide-laced site during egg-laying (**Fig. 4c**). Rather, it increased the proportion of individuals that distributed their eggs over both egg-laying cups (**Fig. 4d**), suggesting that larval exposure increases the likelihood of using skip oviposition, which is defined as visiting (and laying into) several egg-laying sites ^44^. While this is common behaviour in *Aedes* mosquitoes ^45–47^, it is rarely reported in *Anopheles* ^48–50^, which tend to lay all eggs into a single site. A possible reason is that larval exposure to permethrin damages the odour sensors, so that the mosquitoes can no longer distinguish laying sites. Alternatively, it could indicate bet-hedging behaviour, where exposed females decide to lay into several sites to ensure that at least some of their offspring will survive, at risk of losing the offspring laid into the wrong site.

In conclusion, we showed that sublethal larval exposure to insecticides alters, in the long-term, the sensitivity of adult mosquitoes, challenging studies in which pyrethroids appear to remain repellent for resistant mosquitoes ^51^ and reduce the blood-meal size of mosquitoes facing treated nets ^34^. These findings raise important epidemiological concerns, as sublethal exposure increases mosquito motivation to bite and appears to reduce the effectiveness of treated nets against resistant strains like RSP. Since blood-meal size is a crucial parameter influencing malaria transmission potential, our study underscores the issue of insecticide runoff and leaching. It highlights the potential consequences of overusing insecticides and emphasizes the need to seek new, non-insecticide-based control methods.

## Material and Methods

### Experimental system

We used two colonies of the mosquito *An. gambiae* (s.s) – Kisumu a permethrin-sensitive strain ^52^ and RSP a permethrin-resistant strain ^53,54^ – to compare the effects of a prolonged larval exposure to a sublethal dose of the insecticide permethrin on insecticide resistance and various behaviours of adult mosquitoes. The different traits measured were considered in separate experiments that used the same protocol to rear larvae and the same concentration of permethrin (as described below). The resistance of RSP is a product of a *kdr* sodium channel mutation allele, increased cytochrome P450 activity and elevated beta-esterase activity. L4 larvae are exposed to 0.5 mg/L for 24 hours every three generations to maintain permethrin resistance. We maintained the two colonies in three cages of overlapping generations (with an age difference of one week) at a density of about 600 individuals per cage and standard lab conditions (26 ± 1ºC, 70 ± 5% relative humidity and 12 h light/dark) for several years before the experiments.

### Determination of the sublethal doses

Since permethrin has a high absorption in plastic material and a low solubility in water ^55^, all experiments were performed with glass material and the pyrethroid was dissolved in ethanol. Therefore, to control for a potential negative effect of the ethanol, our controls contained the same concentration of ethanol as their respective permethrin treatments. An initial solution was made from solid permethrin (Sigma-Aldrich, Inc., St. Louis, Missouri) dissolved in pure ethanol to obtain a 1mg/L solution, which was used to form the diluted solutions at the concentrations described below.

Larvae were exposed to either ethanol or ethanol supplemented with permethrin throughout their aquatic stage (**Fig. 5a**). The sublethal dose was defined as the concentration for which mortality did not differ significantly (in our laboratory conditions) from the control solution. For Kisumu, based on the sublethal dose previously reported ^17^, we tested three concentrations (0.05, 0.15, 0.45 µg/l) and found that 0.05 µg/l was the sublethal dose. For RSP, we tested 15 concentrations from 0.05 µg/l (the sublethal dose of Kisumu) to 100 µg/l and found that 0.75 µg/l was the sublethal dose. We used these doses in the experiments.

### Mosquito rearing and maintenance

Freshly hatched (0-3h old) Kisumu and RSP larvae were put individually into 60x12mm glass Petri dishes containing 4 ml of permethrin or ethanol solution (**Fig. 5b**). The larvae were fed daily with Tetramin Baby® fish food according to their age (0.04, 0.06, 0.08, 0.16, 0.32 and 0.6 mg/larva respectively for ages 0, 1, 2, 3, 4 and 5 or older ^29^ and some developmental traits were assessed (**Fig. S1**). Pupae were moved to 150 ml cups (5 cm Ø x 10 cm) containing about 50 ml of deionized water and then placed in cages according to their treatment, where they emerged as adults. Adults were fed with 10% sucrose solution placed inside a 150 ml cup (5 cm Ø x 10 cm) that contained a 55 mm Ø Petri dish (to keep the mosquitoes from drowning) on the surface and two 10 x 7 cm rectangular filter paper (to allow mosquitoes to drink the solution). This cup was replaced weekly.

### Insecticide resistance

We tested (1) if prolonged larval exposure to a sublethal dose of permethrin influenced the insecticide resistance of adult mosquitoes, and if this effect was affected by (2) the mosquito strain or (3) the duration of exposure as adults (with two treatment, 30 and 60 min). Males were removed from the cages within the first 24 hours following their emergence to prevent any mating with females used in the experiment. For each treatment, five replicate groups of around 15-20 females were placed into the resting part of WHO test tubes ^56^. After two min of acclimatization, the mosquitoes were gently blown into the exposure part of the tube containing a 0.75% permethrin-impregnated paper (WHO standard paper) for 30 or 60 minutes. During the exposure, we monitored the proportion of individuals that were knockdown at 5, 10, 15, 20, 25 and 30 minutes for all treatments but also at 45 and 60 minutes for the treatments that were exposed for 60 minutes (**Fig. 5c**). After exposure, mosquitoes were gently transferred to the resting part of WHO test tubes with constant access to a 10% sucrose solution and mortality 24 hours post exposure was recorded.

### Choice of protected or unprotected host

We tested (1) if prolonged larval exposure to a sublethal dose of permethrin influenced the choice of protected or unprotected host, and if this effect was affected by (2) the mosquito strain or (3) the time they spent mating. Pupae were collected and put in cages with continuous access to a 10% sucrose solution and upon emergence, males from each treatment were removed from these cages and untreated males were added. Then they were left to mate for 5 or 15 days and we assessed their host-seeking behaviour. The experiment was done with a two-way choice apparatus ^27^, where mosquitoes were able to choose a way in a plastic cage and blood feed either through a permethrin-treated net (Olyset Plus®) or an untreated net (Pharmavoyage® Trek). A ventilator guided the odours (at a speed of about 20 cm/s) from the two lateral cages into a central cage, where the mosquitoes were priorly placed to acclimatize. The side of the insecticide was alternated among tests so that any side preference was avoided (**Fig. 5d**). Females were tested in a treatment per group of approximately 20 individuals, put in a cup and placed in the central cage for 2 minutes for acclimatization. Then, we released them and they had 10 minutes to make a choice. For each mosquito, we noted if they left the central cage, and for the ones that did, the side they chose (with or without permethrin) and if they had a blood meal. Finally, blood-fed mosquitoes were then frozen at − 20 °C for further analysis of the blood meal size.

### Motivation and ability to bite through a permethrin-treated net

We tested (1) if prolonged larval exposure to a sublethal dose of permethrin influenced the motivation to take a blood meal of adult mosquitoes, and if this effect was affected by (2) the mosquito strain or (3) the type of net through which mosquitoes could take a blood meal. Pupae were collected and put in cages with continuous access to a 10% sucrose solution and upon emergence, males from each treatment were removed from these cages and untreated males were added. Then they were left to mate for seven days and on the 7th day, we allowed them to blood-fed for five minutes on TGZ’s arm (**Fig. 5e**). Half of the individuals could blood feed through a permethrin-treated net (Olyset Plus®) and the other half through an untreated net (Pharmavoyage® Trek). We measured the proportion of mosquitoes that took a blood meal on each of the two nets and the mosquitoes were then frozen at − 20 °C for further analysis of the blood meal size.

### Choice of egg-laying site

We tested (1) if a prolonged larval exposure to a sublethal dose of permethrin influenced the egg-laying preference of adult mosquitoes, and if this effect was affected by (2) the mosquito strain. In addition to the larvae reared as described above, some extra larvae were reared in deionized water and upon emergence, males were kept to mate with treated females. This was done to prevent any effect due to treated males. Therefore, at the end of the larval stage, pupae were collected and put in cages with continuous access to a 10% sucrose solution and upon emergence males from each treatment were removed from these cages and untreated males were added. Then they were left to mate for seven days.

On the 7th day, the mosquitoes were blood-fed on TGZ’s arm for five minutes. Two days later (to avoid mortality given the female’s fragility post blood meal) they were individually placed into cages, and given the choice to lay their eggs into a cup (5 cm Ø x 10 cm) containing about 50 ml of deionized water or into one with 50 ml of permethrin solution (**Fig. 5f**). The concentration of the permethrin solution was twice the sublethal dose for each strain (i.e. 0.10 μg/L and 0.0015mg/L for Kisumu and RSP, respectively). The mosquitoes were kept in the cages for two days, with continuous access to a 10% sucrose solution. Then, all cups were collected, filtrated and a picture was taken of the eggs. Eggs were counted using the software ImageJ v1.54 ^57,58^.

### Quantification of blood meal size

Blood meal size was quantified as previously described ^59^. We added a 5 mm Stainless steel bead and 0.5 mL of Drabkin’s reagent to the standards and the samples and grounded them until disintegration using a Qiagen TissueLyser LT at a frequency of 30 hz for two minutes. Then, after a 20 minutes incubation at room temperature (25º C), 0.5 mL of pure chloroform solution was added. Next, we centrifugated at 5600 rpm for 5 minutes and for all samples the supernatant (around 430μL), containing HiCN, was placed in a new 2 mL Eppendorf tube. Finally, after vortexing the supernatant, 2 x 200 μL was placed in a microplate with two technical replicates for the reading of each mosquito and standards. The absorbance was read at 550 nm using a microplate spectrophotometer (SpectraMax i3x plate reader). To be as precise as possible, the standard curves were also made with TGZ’s blood.

### Statistical analysis

All analyses were done with the R software ^60^ version 4.3.0, using the packages “DHARMa” ^61^, “car” ^62^, “lme4” ^63^, “emmeans” ^64^, “multcomp” ^65^ and “survival” ^66^. Significance was assessed with the “Anova” function of the “car” package ^62^. We used a type III ANOVA in the case of a significant interaction and a type II ANOVA otherwise. When relevant, we performed post-hoc multiple comparisons with the package “emmeans”, using the default Tukey adjustment.

### Determination of sublethal doses of permethrin

The sublethal dose was analysed with a generalized linear model with a binomial distribution of errors, where the response variable was the proportion of dead larvae, and the explanatory variables were larval exposure to permethrin and the concentration.

### Insecticide resistance

Knockdown rate was analyzed with a Cox’s proportional hazard model from the survival library ^66^, where the response variable was age at knockdown, the explanatory variables were strain, larval exposure to permethrin, and duration of adults’ exposure and the tube in which the mosquito were exposed as a random factor. The proportional hazard ratio assumption was tested using the function cox.zph from the survival library ^66^. Tubes were tested as a random factor beforehand.

Adult mortality 24 hours after exposure was analysed with a generalized linear model with a binomial distribution of errors, where the response variable was the proportion of dead individuals and the explanatory variables were strain, larval exposure to permethrin and duration of adults’ exposure.

### Choice of protected or unprotected host

Individuals (1) leaving the central cage, (2) choosing the untreated net side and (3) where individuals took a blood-meal were analysed with a generalized linear model with a binomial distribution of errors, where the response variables were the proportion of individuals that (1) left the central cage, (2) went to the side with the untreated net and (3) that took a blood-meal, and the explanatory variables were strain, larval exposure to permethrin and for how long they were allowed to mate.

Blood-meal size was analysed with a linear model with a Gaussian distribution of errors, where the response variable was the individual quantity of haemoglobin in μg/mosquito and the explanatory variables were strain, larval exposure to permethrin and for how long they were allowed to mate.

### Motivation and ability to bite through a permethrin-treated net

Individual motivation to take a blood-meal was analysed with a generalized linear model with a binomial distribution of errors, where the response variable was the proportion of individuals that took a blood-meal and the explanatory variables were strain, larval exposure to permethrin and the type of net they could bite through.

Blood-meal size was analysed with a linear model with a Gaussian distribution of errors, where the response variable was the individual quantity of haemoglobin in μg/mosquito, and the explanatory variables were strain, larval exposure to permethrin and the type of net they could bite through.

### Choice of egg-laying site

Individuals (1) not laying any egg, (2) laying in both egg-layers and (3) choosing the insecticide-laced egg-layer were analysed with three generalized linear models with a binomial distribution of errors, where the response variables were (1) the proportion of individuals that did not laid any egg, or (for mosquitoes that laid eggs) (2) the proportion of individuals laid into both egg-laying cups or (3) the proportion of eggs on the insecticide-laced egg-laying cup, and the explanatory variables were strain and larval exposure to permethrin.

The number of eggs was analysed with a linear model with a Gaussian distribution of errors, where the response variable was individual egg count, and the explanatory variables were strain and larval exposure to permethrin.

## Supporting information

Supplementary information for " Sublethal insecticide exposure of larvae affects the blood-feeding behaviour of adult mosquitoes "

## Acknowledgements

We thank Luís M. Silva for his advice and technical support and all the students that were involved in this project: Fanchon Cachin, Camille Lattion, Quentin Clot, Vanessa Herren, Rhéanne Udriot, Léonard Nigra and Melvin Chavanne. TGZ, GA and the project were supported by SNF grant 310030_192786.

## Author contributions

TGZ and JCK conceived the overall idea. TGZ designed the experiments. TGZ and GA collected the data. TGZ and GA analysed the data and wrote the first draft of the manuscript. All authors contributed critically to the drafts.

## Data availability

All data generated or analysed during this study are included as Supplementary Information files.

## Additional Information

The authors declare no competing interests.

## Notes

### Competing Interest Statement

The authors have declared no competing interest.

