## Supplementary information for " Sublethal insecticide exposure of larvae affects the blood-feeding behaviour of adult mosquitoes " for "Sublethal insecticide exposure of larvae affects the blood-feeding behaviour of adult mosquitoes"

**Figure S1.** Developmental traits

**Table S1.** Insecticide resistance

**Table S2.** Choice of protected or unprotected host

**Table S3.** Motivation and ability to bite through a permethrin-treated net

**Table S4.** Egg-laying preference

**Table S5.** Developmental traits

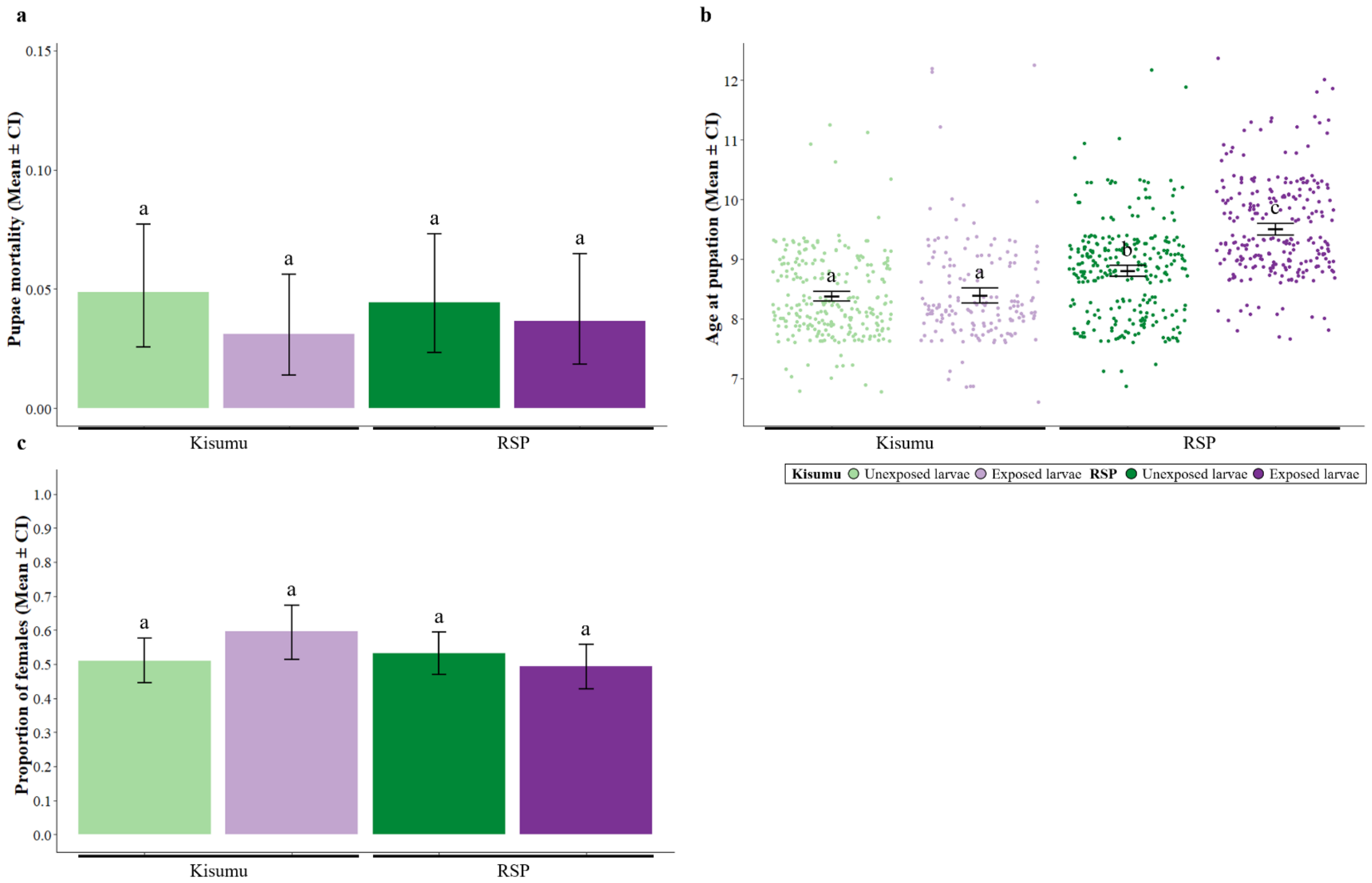


**Figure S1. Developmental traits.** The proportion of Kisumu and RSP individuals, exposed or unexposed to permethrin during the larval stage that died **(a)** as pupae, **(b)** their age at pupation and **(c)** their sex ratio (as the proportion of females). Letters indicate statistically significant differences from multiple comparisons. Pupae mortality was analysed with a generalized linear model with a binomial distribution of errors, where the response variable was the proportion of dead pupae and the explanatory variables were strain and larval exposure to permethrin. Age at pupation was analysed with a linear model with a Gaussian distribution of errors, where the response variable was age at pupation and the explanatory variables were strain, larval exposure to permethrin and sex. Sex ratio was analysed with a generalized linear model with a binomial distribution of errors, where the response variable was the sex of the mosquito and the explanatory variables were strain, larval exposure to permethrin and age at pupation. For details on the sample size see **Supplementary Table S5.2**.

Pupae mortality was 4% in Kisumu-mosquitoes and RSP-mosquitoes (0.04 for both, $\chi^2 = 0.00$, df = 1, p = 0.995), and was not affected by larval exposure to permethrin ($\chi^2 = 0.82$, df = 1, p = 0.362) or the interaction between larval exposure and strain ($\chi^2 = 0.14$, df = 1, p = 0.703, **Fig. S1a**).

RSP-mosquitoes took more than half a day longer to pupate than Kisumu-mosquitoes (9.1 days vs 8.38 days, $F_{1,879} = 30.22$, df = 1, $p < 0.001$), and females took longer than males (8.88 days vs 8.63 days, $F_{1,879} = 8.93$, df = 1, p = 0.002). However, we also found an effect of exposure to permethrin that was only evident for RSP-mosquitoes, with exposed individuals pupating later than unexposed (8.32 days unexposed vs 8.36 days exposed for Kisumu-mosquitoes and 8.77 days unexposed vs 9.46 days exposed for RSP-mosquitoes, interaction strain * exposure: $F_{1,879} = 34.05$, df = 1, $p < 0.001$, **Fig. S1b**).

The proportion of females among the adults was higher for Kisumu-mosquitoes than for RSP-mosquitoes (0.54 vs 0.51, $\chi^2 = 11.51$, df = 1, $p < 0.001$), but was not affected by exposure to permethrin (0.52 for unexposed vs 0.53 for exposed, $\chi^2 = 1.26$, df = 1, p = 0.261). However, we found an effect of exposure to permethrin that was only evident for Kisumu-mosquitoes with exposed individuals leading to a higher proportion of females than unexposed (0.51 unexposed vs 0.59 exposed for Kisumu-mosquitoes and 0.53 unexposed vs 0.49 exposed for RSP-mosquitoes, interaction strain * exposure: $\chi^2 = 6.81$, df = 1, p = 0.009, **Fig. S1c**). In addition, the proportion of females also increased with later pupation ($\chi^2 = 41.94$, df = 1, $p < 0.001$). For further details on the statistical analysis see **Supplementary Table S5.1**.

**Table S1.1. Insecticide resistance**

| **Tested effects** | | | |
|---|---|---|---|
| **Model 1a** - Knock-downs | **df** | $\chi^2$ | ***p*** |
| Exposure | 1 | 0.27 | 0.605 |
| Strain | 1 | 531.24 | **<0.001** |
| Exposure time | 1 | 87.45 | **<0.001** |
| Strain : Exposure | 1 | 0.34 | 0.561 |
| Strain : Exposure time | 1 | 20.64 | **<0.001** |
| Exposure : Exposure time | 1 | 14.30 | **<0.001** |
| Strain : Exposure : Exposure time | 1 | 1.01 | 0.315 |
| **Model 1b** - Adult mortality | | | |
| Exposure | 1 | 2.34 | 0.125 |
| Strain | 1 | 180.83 | **<0.001** |
| Exposure time | 1 | 3.09 | 0.078 |
| Strain : Exposure | 1 | 1.78 | 0.180 |
| Strain : Exposure time | 1 | 20.65 | **<0.001** |
| Exposure : Exposure time | 1 | 1.40 | 0.235 |
| Strain : Exposure : Exposure time | 1 | 1.63 | 0.200 |

**Table S1.2. Insecticide resistance – Sample size**

**Sample size**

| | Kisumu | | | | RSP | | | |
|---|---|---|---|---|---|---|---|---|
| | Unexposed | | Exposed | | Unexposed | | Exposed | |
| | 30 min | 60 min | 30 min | 60 min | 30 min | 60 min | 30 min | 60 min |
| | 196 | 177 | 167 | 150 | 121 | 122 | 133 | 131 |

**Table S2.1. Choice of protected or unprotected host**

| Tested effects | | | | |
|---|---|---|---|---|
| **Model 2a** - Probability of leaving the central cage | **df** | | $\chi^2$ | ***p*** |
| Exposure | 1 | | 0.97 | 0.322 |
| Strain | 1 | | 1.41 | 0.234 |
| Mating duration | 1 | | 3.39 | 0.065 |
| Strain : Exposure | 1 | | 3.58 | 0.058 |
| Strain : Mating duration | 1 | | 13.01 | **<0.001** |
| Exposure : Mating duration | 1 | | 0.06 | 0.799 |
| Strain : Exposure : Mating duration | 1 | | 1.10 | 0.293 |
| **Model 2b** - Probability of choosing the normal net side | | | | |
| Exposure | 1 | | 6.08 | **0.013** |
| Strain | 1 | | 4.22 | **0.039** |
| Mating duration | 1 | | 3.88 | **0.048** |
| Strain : Exposure | 1 | | 0.23 | 0.626 |
| Strain : Mating duration | 1 | | 0.62 | 0.428 |
| Exposure : Mating duration | 1 | | 0.01 | 0.907 |
| Strain : Exposure : Mating duration | 1 | | 3.78 | 0.051 |
| **Model 2c** - Probability of taking a BM | | | | |
| Exposure | 1 | | 1.29 | 0.255 |
| Strain | 1 | | 5.19 | **0.022** |
| Mating duration | 1 | | 0.89 | 0.343 |
| Strain : Exposure | 1 | | 4.20 | **0.040** |
| Strain : Mating duration | 1 | | 1.04 | 0.306 |
| Exposure : Mating duration | 1 | | 0.30 | 0.581 |
| Strain : Exposure : Mating duration | 1 | | 2.68 | 0.101 |
| **Model 2d** - BM size | | Sum Sq | F | *p* |
| Exposure | 1 | 79 | 0.06 | 0.804 |
| Strain | 1 | 2081 | 1.61 | 0.204 |
| Mating duration | 1 | 3682 | 2.86 | 0.092 |
| Strain : Exposure | 1 | 216 | 0.16 | 0.682 |
| Strain : Mating duration | 1 | 13176 | 10.25 | **0.001** |
| Exposure : Mating duration | 1 | 473 | 0.36 | 0.544 |
| Strain : Exposure : Mating duration | 1 | 56 | 0.04 | 0.834 |

**Table S2.2. Choice of protected or unprotected host – Sample size**

| Sample size | | | | | | | | |
|---|---|---|---|---|---|---|---|---|
| | **Kisumu** | | | | **RSP** | | | |
| | **Unexposed** | | **Exposed** | | **Unexposed** | | **Exposed** | |
| | **5 days** | **15 days** | **5 days** | **15 days** | **5 days** | **15 days** | **5 days** | **15 days** |
| Start | 52 | 59 | 80 | 92 | 96 | 86 | 104 | 118 |
| After leaving central cage | 15 | 27 | 17 | 36 | 37 | 14 | 54 | 18 |
| After took a BM | 15 | 26 | 16 | 30 | 30 | 12 | 53 | 11 |

**Table S3.1. Motivation and ability to bite through a permethrin-treated net**

| **Tested effects** | | | | |
|---|---|---|---|---|
| **Model 3a** - Probability blood feeding (BF) | | | | |
| Exposure | 1 | | 0.92 | 0.335 |
| Strain | 1 | | 13.84 | **<0.001** |
| Net | 1 | | 0.91 | 0.337 |
| Strain : Exposure | 1 | | 0.96 | 0.326 |
| Strain : Net | 1 | | 5.79 | **0.016** |
| Exposure : Net | 1 | | 4.74 | **0.029** |
| Strain : Exposure : Net | 1 | | 0.003 | 0.954 |
| **Model 3b** - Blood meal (BM) size | **df** | **Sum Sq** | **F** | ***p*** |
| Exposure | 1 | 20130 | 17.93 | **<0.001** |
| Strain | 1 | 3981 | 3.54 | 0.060 |
| Net | 1 | 39 | 0.03 | 0.852 |
| Strain : Exposure | 1 | 6732 | 5.99 | **0.014** |
| Strain : Net | 1 | 392 | 0.34 | 0.554 |
| Exposure : Net | 1 | 2883 | 2.56 | 0.110 |
| Strain : Exposure : Net | 1 | 6781 | 6.04 | **0.014** |

**Table S3.2. Motivation and ability to bite through a permethrin-treated net**

**Sample size**

| | **Kisumu** | | | | **RSP** | | | |
|---|---|---|---|---|---|---|---|---|
| | **Unexposed** | | **Exposed** | | **Unexposed** | | **Exposed** | |
| | **Normal net** | **Perm. net** | **Normal net** | **Perm. net** | **Normal net** | **Perm. net** | **Normal net** | **Perm. net** |
| Start | 74 | 78 | 75 | 73 | 75 | 60 | 77 | 70 |
| After taking a BM | 45 | 36 | 45 | 46 | 30 | 27 | 24 | 24 |

**Table S4.1. Choice of egg-laying site**

| **Tested effects** | | | | |
|---|---|---|---|---|
| **Model 4a** - Probability not laying | **df** | | $\chi^2$ | ***p*** |
| Exposure | 1 | | 0.02 | 0.882 |
| Strain | 1 | | 22.58 | **<0.001** |
| Strain : Exposure | 1 | | 0.76 | 0.380 |
| **Model 4b** - Proportion of eggs in the insecticide-laced site | | | | |
| Exposure | 1 | | 0.06 | 0.802 |
| Strain | 1 | | 1.28 | 0.257 |
| Strain : Exposure | 1 | | 0.55 | 0.455 |
| **Model 4c** - Probability laying in both egg-layers | | | | |
| Exposure | 1 | | 6.05 | **0.013** |
| Strain | 1 | | 3.39 | 0.065 |
| Strain : Exposure | 1 | | 2.53 | 0.111 |
| **Model 4d** - Egg count | **df** | **Sum Sq** | **F** | ***p*** |
| Exposure | 1 | 851 | 0.93 | 0.333 |
| Strain | 1 | 52677 | 58.09 | **<0.001** |
| Strain : Exposure | 1 | 3186 | 3.51 | 0.061 |

**Table S4.2. Choice of egg-laying site – Sample size**

| Sample size | | | | |
|---|---|---|---|---|
| | Kisumu | | RSP | |
| | Unexposed | Exposed | Unexposed | Exposed |
| Start | 90 | 88 | 89 | 86 |
| Individuals that laid eggs | 83 | 78 | 62 | 62 |

**Table S5.1. Developmental traits**

| **Tested effects** | | | | |
|---|---|---|---|---|
| **Model 5a** - Sublethal doses *Kisumu* | **df** | | $\chi^2$ | ***p*** |
| Exposure | 1 | | 109.14 | **<0.001** |
| Concentration | 1 | | 68.04 | **<0.001** |
| Exposure : Concentration | 1 | | 3.31 | 0.068 |
| **Model 5b** - Sublethal doses *RSP* | | | | |
| Exposure | 1 | | 454.52 | **<0.001** |
| Concentration | 1 | | 1095.77 | **<0.001** |
| Exposure : Concentration | 1 | | 127.35 | **<0.001** |
| **Model 5c** - Pupal mortality | | | | |
| Exposure | 1 | | 0.82 | 0.362 |
| Strain | 1 | | 0.00 | 0.995 |
| Exposure : Strain | 1 | | 0.14 | 0.703 |
| **Model 5d** - Proportion of females | | | | |
| Exposure | 1 | | 1.26 | 0.261 |
| Strain | 1 | | 11.51 | **<0.001** |
| Age at pupation | 1 | | 41.94 | **<0.001** |
| Strain : Exposure | 1 | | 6.81 | **0.009** |
| Strain : Age at pupation | 1 | | 0.19 | 0.657 |
| Exposure : Age at pupation | 1 | | 0.37 | 0.539 |
| Strain : Age at pupation : Exposure | 1 | | 2.34 | 0.125 |
| **Model 5e** - Age at pupation | **df** | **Sum Sq** | **F** | ***p*** |
| Exposure | 1 | 0 | 0.009 | 0.923 |
| Strain | 1 | 14.2 | 30.22 | **<0.001** |
| Sex | 1 | 4.2 | 8.93 | **0.002** |
| Strain : Exposure | 1 | 16.0 | 34.05 | **<0.001** |
| Strain : Sex | 1 | 0.1 | 0.11 | 0.738 |
| Exposure : Sex | 1 | 0.1 | 0.31 | 0.574 |
| Strain : Exposure : Sex | 1 | 0.3 | 0.67 | 0.412 |

**Table S5.2. Developmental traits – Sample size**

| Sample size | | | | |
|---|---|---|---|---|
| | **Kisumu** | | **RSP** | |
| | **Unexposed** | **Exposed** | **Unexposed** | **Exposed** |
| Sublethal concentrations | 100/concentration | 100/concentration | 50/concentration | 50/concentration |
| Developmental traits | 300 | 300 | 300 | 300 |
